# 3D Bioprinting of Graphene Oxide-Incorporated Cell-laden Bone Mimicking Scaffolds for Promoting Scaffold Fidelity, Osteogenic Differentiation and Mineralization

**DOI:** 10.1101/2020.08.14.251074

**Authors:** Jianhua Zhang, Hande Eyisoylu, Xiao-Hua Qin, Marina Rubert, Ralph Müller

## Abstract

Bioprinting is a promising technique for facilitating the fabrication of engineered bone tissues for patient-specific defect repair and for developing *in vitro* tissue/organ models for *ex vivo* tests. However, polymer-based ink materials often result in insufficient mechanical strength, low scaffold fidelity and loss of osteogenesis induction because of the intrinsic swelling/shrinking and bioinert properties of most polymeric hydrogels. In this work, we developed a novel human mesenchymal stem cell (hMSC)-laden graphene oxide (GO)/alginate/gelatin composite bioink to form 3D bone mimicking scaffolds. Our results showed that the GO composite bioinks with higher GO concentrations improved the bioprintability, scaffold fidelity, compressive modulus and cell viability. The higher GO concentration increased the cell body size and DNA content. The 1GO group had the highest osteogenic differentiation of hMSC with the upregulation of osteogenic-related gene expression at day 42. To mimic critical-sized calvarial bone defects in mice, 3D cell-laden GO defect scaffolds with complex geometries were successfully bioprinted. 1GO maintained the best scaffold fidelity and had the highest mineral volume after culturing in the bioreactor for 42 days. Finally, the 1GO bioink has been demonstrated great potential for 3D bioprinting in applications of bone model and bone tissue engineering.

## 1. Introduction

Bone fractures and bone diseases, such as osteoporosis, are a significant source of patient morbidity and place a staggering economic burden on the healthcare system. Developing treatments for bone diseases and fracture healing requires an understanding of the molecular mechanisms and mineral formation in bone developmental biology. 3D bioprinting techniques have shown promise for mimicking the structure of different tissues and organs through the precise placement of living cells and biomaterials in a layer-by-layer fashion ^[1]^. 3D bioprinting has opened a new perspective for bone defect bioprinting using patient-specific medical images ^[2]^ and for developing *in vitro* tissue/organ models for *ex vivo* tests ^[3]^. One challenge in bone bioprinting remains supporting high cell viability after bioprinting and inducing extracellular matrix (ECM) mineralization *in vitro*. Different hydrogel materials (alginate, gelatin, gelatin methacryloyl, fibrinogen, etc.) and cell types have been successfully bioprinted for bone tissue engineering ^[4]^. Among them, alginate-gelatin blend inks have been widely used for 3D bioprinting due to their biocompatibility, low cost and instant gelation at room temperature ^[5]^. The thermosensitive properties of gelatin facilitate the initial stability of 3D bioprinted constructs ^[6]^. Alginate polymer chains are crosslinked with Ca^2+^ after bioprinting to provide long-term mechanical strength. Our previous studies have shown that low-density alginate/gelatin (0.8% alg) blend bioinks can maintain high cell viability (> 85%) and induce the osteogenic differentiation of human mesenchymal stem cells (hMSC) ^[7]^. Significantly more mineralized tissue was formed in low-density (0.8% alg) than high-density (1.8%) alginate/gelatin scaffolds (43.5 ± 7.1 mm^3^ *vs* 22.6 ± 6.0 mm^3^) after culturing in osteogenic media for 42 days. However, low-density alginate/gelatin ink solutions often result in moderate printability, insufficient mechanical strength, and low scaffold fidelity because of the intrinsic swelling and/or shrinking properties of alginate and gelatin. The scaffold fidelity and mechanical properties could be improved with higher density bioinks, such as by increasing alginate concentration, while increased viscosity suppressed cell viability and cell morphology ^[7]^. Thus, the ability to design low-density alginate/gelatin-based bioinks with improved mechanical properties, scaffold fidelity and cell functions remains a challenge in 3D bioprinting for engineering bone tissues ^[8]^.

Organic/inorganic composite ink formulations have been studied to obtain the desired mechanical properties and gel behaviors for 3D bioprinting ^[9]^. The most common inorganic particles added to hydrogels to prepare ink solutions are silica ^[10]^, laponite ^[11]^, hydroxyapatite ^[9b]^, gold and silver nanoparticles ^[12]^. However, these inorganic nanoparticles increase shear stress during the bioprinting processes, which may suppress cell viability and affect cell functions ^[8]^. Nanoparticles may limit the diffusion of crosslinking reagents, resulting in inhomogeneous crosslinking of the relevant large 3D constructs ^[13]^. Graphene oxide (GO), readily prepared from the oxidation of graphite, is an exciting nanomaterial for tissue engineering and regenerative medicine. GO has an atomically thin sheet with a large surface area and abundant hydrophilic functional groups (e.g., hydroxyl, epoxide and carbonyl) that allow a wide range of chemical modifications, and GO exhibits excellent absorption properties ^[14]^. 2D studies have shown that GO improves hMSC adhesiveness and proliferation ^[15]^, cell growth ^[16]^ and osteogenic differentiation ^[16c]^. In 3D studies, many researchers have incorporated GO into hydrogel scaffolds, such as alginate ^[17]^, collagen ^[18]^ and gelatin methacrylate (GelMA) ^[19]^, and polymer chains were crosslinked to form a 3D GO hybrid scaffold. For example, Su et al. ^[19]^ incorporated GO into GelMA for the creation of cell-laden graphene-embedding hydrogels and investigated the cellular responses in a 3D microenvironment. They demonstrated that GO-GelMA hybrid hydrogels supported the spreading and alignment of fibroblasts with improved viability and proliferation. Marrella et al. ^[17]^ confirmed an absence of cytotoxicity of suspended GO flakes and improved viability of fibroblasts encapsulated in 3D GO/alginate hydrogels. Because of the negative surface charge and excellent absorbing properties of GO ^[20]^, we hypothesized that the incorporation of GO into the alginate/gelatin ink system improves scaffold fidelity due to the attractive electrostatic forces between GO and Ca^2+^. To our knowledge, the influence of GO incorporation in 3D bioprinted cell-laden scaffolds on osteogenic differentiation and mineral formation has not yet been investigated.

In this study, four different GO composite bioinks were prepared by mixing hMSC into an alginate/gelatin (0.8%/4.1%, w/v) solution containing different GO concentrations (0, 0.5, 1, 2 mg/ml). The influence of GO on bioprintability was first investigated by characterizing the shear thinning and shear recovery properties. The scaffold fidelity and mechanical properties of these varying GO composite scaffolds were studied by analyzing the changes in scaffold morphology and the compressive modulus on day 1, day 7 and day 42. To evaluate the influence of GO on cell behavior and function, we studied cell viability, cell morphology, DNA content, alkaline phosphatase (ALP) activity and osteogenic-related gene expression in a systematic fashion. 3D bioprinting offers exciting prospects for constructing 3D tissue/organ models, as it enables the reproducible, automated production of complex living tissues ^[3]^. The 3D bioprinted tissue/organ model may prove useful as an *ex vivo* model for reducing animal usage and screening novel compounds or predicting toxicity. Here, we bioprinted 3D cell-laden scaffolds with complex geometries to mimic critical-sized calvarial defects in mice. The influence of GO incorporation on bone-like tissue formation in 3D bioprinted cell-laden defect scaffolds was investigated using *in situ* micro-computed tomography (micro-CT) monitoring in 42 days of cell culture.

## 2. Results and discussion

### 2.1 GO characterization, bioink preparation

Commercially available GO was produced through oxidation/exfoliation of graphite powder via the modified Hummer’s method. GO nanoflakes displayed a lateral size of >15 µm and a thickness of 0.8-1.2 nm, according to the manufacturer’s information. The morphology of GO was characterized by TEM analysis (Figure 1A), which showed that the GO was single-layer materials. The exfoliated GO nanosheets were readily dispersed in deionized water with ultrasonic treatment and formed a suspension that was stable for several months with no precipitation. Photograph and optical microscopy analysis (Figure 1B C) showed that the 5 mg/ml GO dispersion in water solution is a homogeneous and stable suspension without aggregation. As GO flasks are brown, the GO flask conjugation to alginate/gelatin chains changed the hydrogel color from white to brown to black according to the GO concentration, as shown in the photograph (Figure 1D). To confirm the dispersion of GOs within the hydrogel solutions, optical microscopy analysis of the GO/alginate/gelatin composite solution was performed. No evidence of aggregation was observed, and a homogeneous distribution of GO flakes throughout the volume was revealed. When the GO concentration was increased from 0.5 to 2 mg/ml in the composite hydrogels, more GO flakes could be observed under the optical microscopy images (Figure 1E-H). Our result is similar to previous findings reported by Guo and Scaglione et al ^[17, 21]^, who demonstrated that the negative ions in both alginate and GO allow the formation of a homogeneous and stable solution.

**Figure 1.**
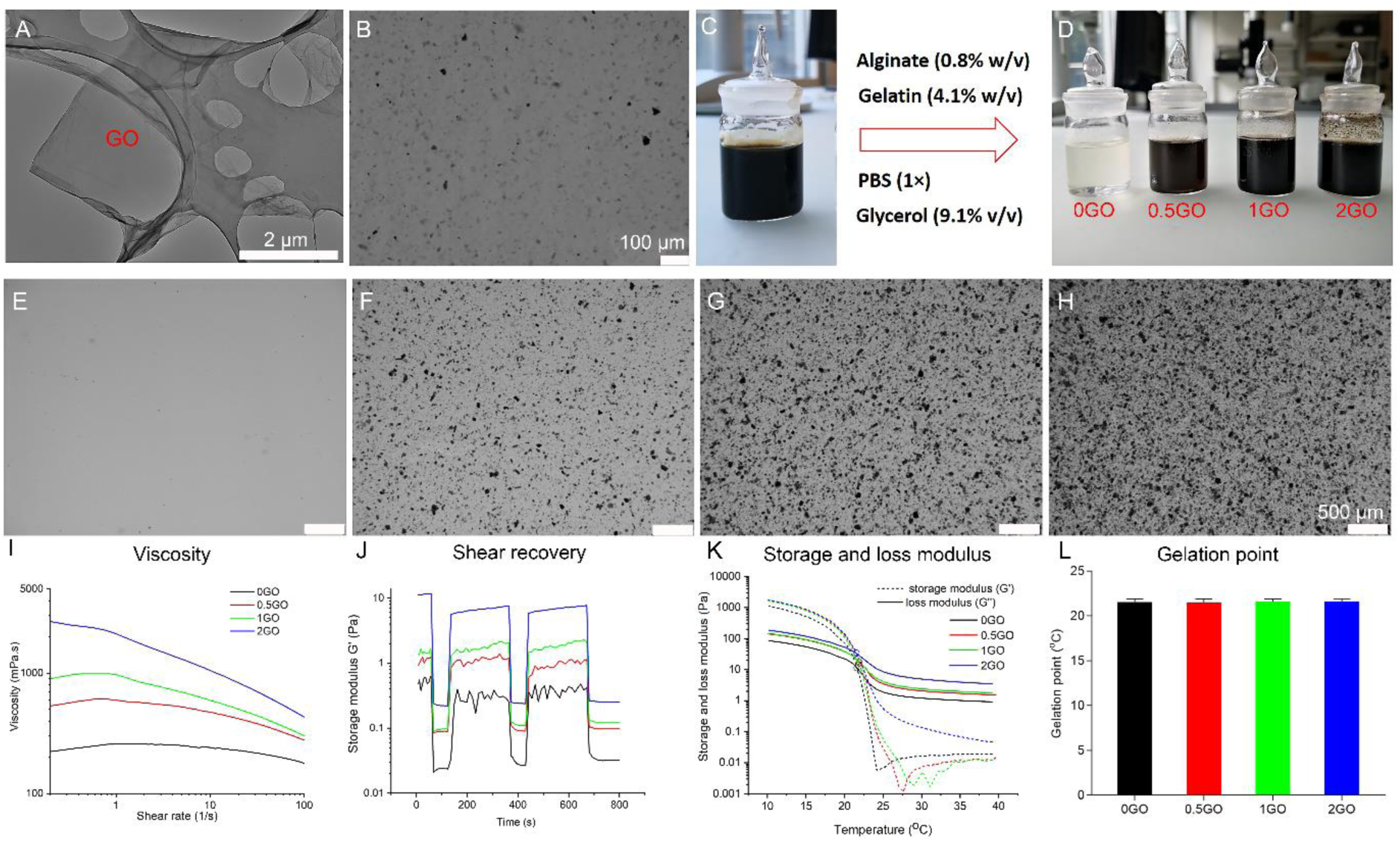
(A) TEM image of graphene oxide (GO). (B) Optical microscopy image and (C) photograph of the 5 mg/ml GO dispersion in water. (D) Photograph of 0GO, 0.5GO, 1GO and 2GO hydrogel solutions. (E-H) Optical microscopy images of 0GO (E), 0.5GO (F), 1GO (G) and 2GO (H). (I-L) Rheological characterization of the ink solutions, (I) Viscosity, (J) Shear recovery, (K) Storage modulus and loss modulus change during cooling from 40 °C to 10 °C, (L) Average gelation point of the four different GO inks (n=3).

### 2.2 Rheological properties

The rheological properties of the hydrogels with varied GO contents were measured to determine the shear behavior and shear recovery, two of the most important predictors of bioink printability. All of the bioink compositions showed shear thinning behavior, while the higher GO content group showed better shear thinning properties (Figure 1I). During shear thinning, the coiled polymer chains align and disentangle at higher shear rates, requiring less extrusion force to deposit the bioink, which is beneficial for cell survival. Shear recovery relates to the ink’s resistance to flow after printing, which ensures high fidelity of the printed structure. The shear recovery curves (Figure 1J) illustrate that incorporating GO into the alginate/gelatin composite hydrogel increases shear recovery. Shear recovery after the first shear sequence was 79.55% in 0.5GO, 97.09% in 1GO and 51.08% in 2GO of the original storage modulus. At the same time, the bioink without GO recovered to only 37.39%. Furthermore, because of the thermosensitive properties of gelatin, forming a stable 3D bioprinted GO/alginate/gelatin construct required decreasing the temperature from 37 °C to 10 °C. Representative curves of the viscoelastic moduli of the different hydrogel groups are shown in Figure 1K. The storage and loss moduli of the GO/alginate/gelatin hydrogel significantly increased during the gelation process, indicating an increase in viscosity and bioink stability. The influence of the GO concentration on the gelation point is shown in Figure 1L. The gelation points for the 0GO, 0.5GO, 1GO and 2GO groups were 21.6 ± 0.3 °C, 21.5 ± 0.3 °C, 21.6 ± 0.2 °C and 21.6 ± 0.1 °C, respectively. There was no statistically significant difference in the gelation point between hydrogels with different GO concentrations. Ma et al. ^[22]^ showed that the gelation point is mostly related to the gelatin concentration in gelatin-based bioink composition.

### 2.3 Scaffold structure

Hydrogels are well known for their swelling behavior, which is an essential consideration in predicting the functioning of the printed geometry and achieving an accurate spatial distribution of material and cells ^[23]^. We evaluated the morphology of 3D bioprinted acellular (Figure 2) and cell-laden (Figure 3) GO composite scaffolds. Light microscopy images showed that 3D acellular and cell-laden GO scaffolds were successfully bioprinted with regular and interconnected macroporous structures. Figure 2 shows that acellular scaffolds had different overall shrinkage compared with the scaffold model (10 × 10 mm) depending on the GO concentration. Increasing the GO concentration showed higher scaffold area fidelity with less scaffold dimension reduction. As shown in Figure 2E at day 7, the acellular scaffold overall area of 0GO was 65.36 ± 3.87 mm^2^, which was significantly lower than 89.60 ± 6.31 mm^2^, 92.39 ± 6.68 mm^2^, 96.08 ± 2.75 mm^2^ obtained for 0.5GO, 1GO and 2GO, respectively. The pore size and pore area of 0GO were 0.58 ± 0.02 mm and 0.28 ± 0.01 mm^2^, respectively, which were not significantly different from those of the GO groups (Figure 2 F G). However, the filament diameters of 0.5GO, 1GO and 2GO were 0.33 ± 0.04, 0.32 ± 0.02, and 0.36 ± 0.01 mm, respectively, which were significantly higher than 0.23 ± 0.02 mm for 0GO at day 7 (Figure 2H).

**Figure 2.**
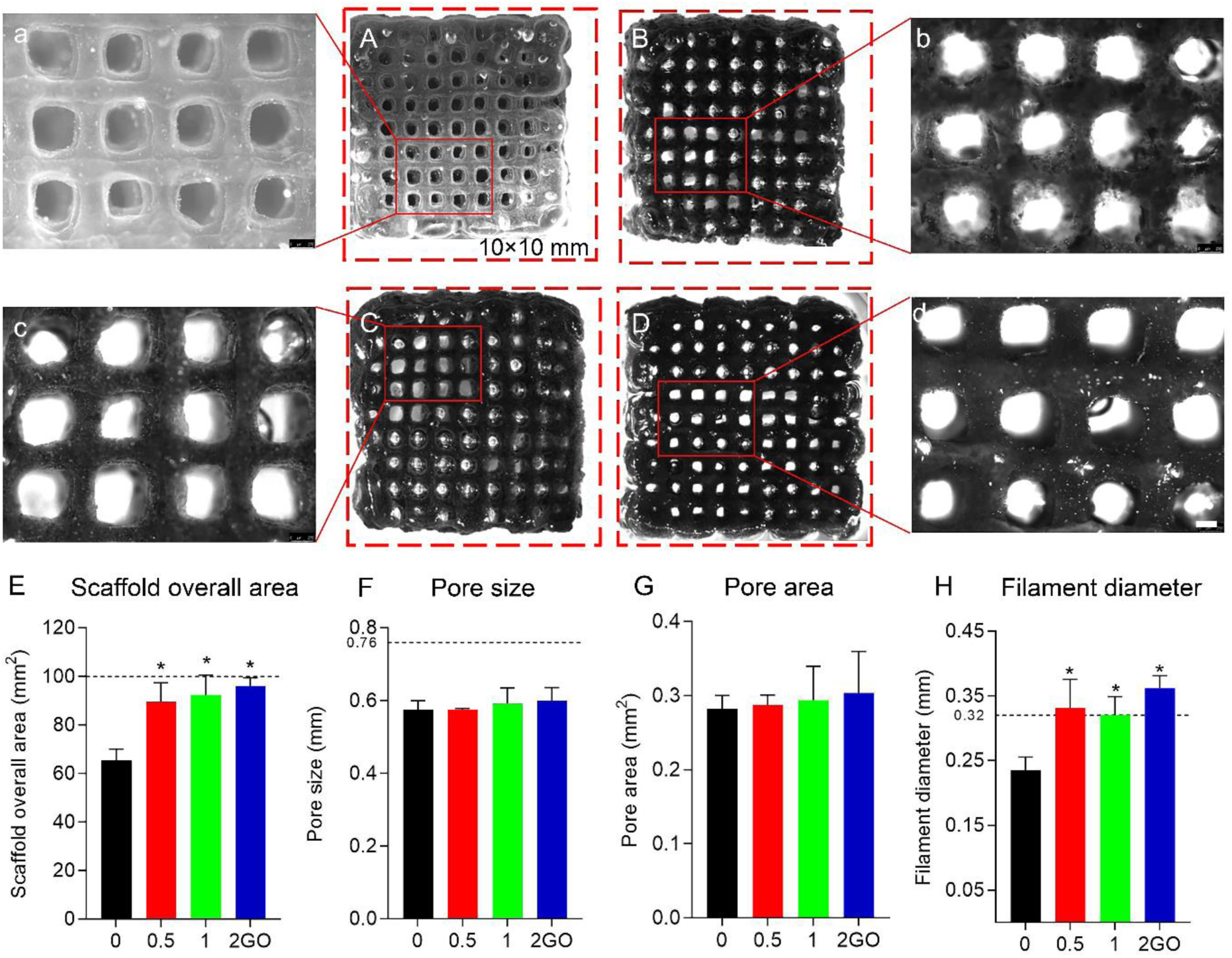
(A-D) (a-d) Light microscopy images of the 3D bioprinted acellular scaffolds cultured in the cell culture media for 7 days. (A, a) 0GO, (B, b) 0.5GO, (C, c) 1GO and (D, d) 2GO. (A-D) Full scaffold images, (a-d) High magnification images of the solid rectangle in the scaffolds, the dimension of the dashed red-line rectangle is 10 × 10 mm. (E-H) Scaffold parameter of the acellular scaffolds at day 7 (n = 3), (E) Scaffold overall area, (F) Pore size, (G) Pore area and (H) Filament diameter. Dashed black lines represent the values of the designed scaffold model. * P < 0.05 was compared to the 0GO group.

**Figure 3.**
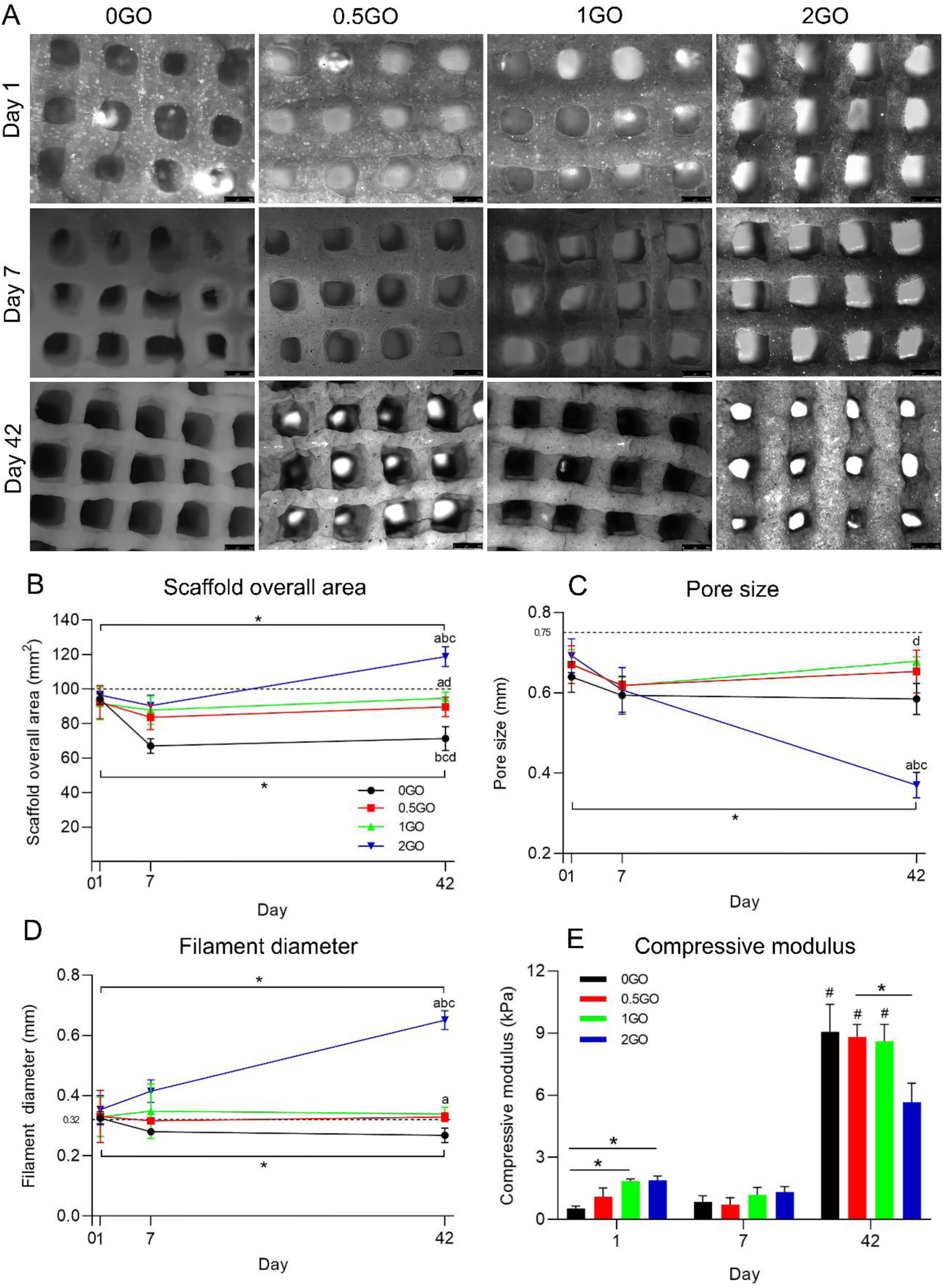
(A) Light microscopy images of the 3D bioprinted cell-laden GO scaffolds cultured in the osteogenic media for 1 day, 7 days and 42 days. Scaffold parameter changes of the 3D bioprinted cell-laden GO scaffolds: (B) Scaffold overall area, (C) Pore size and (D) Filament diameter. ^a b c d^ represent significant differences (P < 0.05) compared to 0GO, 0.5GO, 1GO and 2GO group, respectively, within the same time point. * P < 0.05, within the same group between day 42 and day 1. Data are shown as mean ± standard deviation (n = 3), dashed black lines represent the values of the designed scaffold model. (E) Compressive moduli of the 3D bioprinted cell-laden GO scaffolds after culturing in the osteogenic media for 1 day, 7 days and 42 days. * P < 0.05, and # P < 0.05 between day 42 and day 1, between day 42 and day 7.

Similar changes in scaffold morphology as observed in the acellular scaffolds were found in the 3D bioprinted cell-laden scaffolds after culture in the osteogenic cell culture media for 1 day and 7 days (Figure 3A). 3D cell-laden scaffolds with higher GO content improved the scaffold morphology with less shrinkage over 7 days. However, after long-term culture in osteogenic media for 42 days, the 2GO group exhibited prominent swelling, and no significant differences were detected among the 0GO, 0.5GO and 1GO groups from day 7 to day 42. Pore size, filament diameter and overall scaffold area of the 2GO group significantly differed between day 1 and 42; these parameters in the 2GO group also significantly differed compared with those of the 0GO, 0.5GO and 1GO groups at day 42 (Figure 3 B-D). Over 42 days cultured in the osteogenic media, the 0GO scaffolds were shrinkage and had the lower scaffold overall area and filament diameter compared to 0.5GO, 1GO and 2GO. The best candidate for maintaining scaffold morphology compared with the scaffold design model was the scaffold of the 1GO group.

The shrinkage behavior of 0GO was similar to that observed in our previously published study when the scaffold was cultured in cell culture media ^[7]^. James et al. ^[24]^ and Ma et al. ^[22]^ reported that ionically crosslinked alginate gel will lose its stability due to the loss of crosslinking calcium ions when in the presence of calcium chelators (e.g., phosphates), monovalent ions (e.g., K^+^, Na^+^, etc.), and non-crosslinking divalent ions (e.g., Mg^2+^). However, alginate gels could remain stable in cell culture media because some calcium ions in the media counteract the deprivation effects [25]. Incorporating GO into the 3D bioprinted alginate/gelatin scaffolds may help maintain better scaffold morphology when cultured in the cell culture media in the short term. Some researchers have reported that GO has extraordinary absorption capacity for positive ions ^[26]^, proteins ^[27]^, and small molecule drugs ^[28]^ via electrostatic interactions and π-π stacking. This behavior decreases the loss of crosslinking calcium ions and increases the stability of alginate gel. However, a higher GO concentration will result in the absorption of more hydrophilic proteins and water molecules, which contributes to the swelling behavior in the long term.

### 2.4 Scaffold mechanics

To investigate the effect of incorporating GO on the mechanical properties, unconfined compression tests were performed on 3D bioprinted cell-laden GO scaffolds cultured in osteogenic media for 1, 7 and 42 days. Figure 3E shows the compressive moduli of 3D bioprinted cell-laden scaffolds with a range of GO concentrations (0, 0.5, 1, 2 mg/ml). The incorporation of GO improved the mechanical properties of the 3D bioprinted cell-laden scaffolds on day 1. The compressive moduli of the 1GO and 2GO scaffolds were 1.58 ± 0.28 kPa and 1.63 ± 0.29 kPa, respectively, which were significantly higher than that of 0GO (0.69 ± 0.24 kPa) on day 1. Meanwhile, the compressive moduli of the 0GO, 0.5GO, and 1GO scaffolds were significantly higher than that of the 2GO scaffolds on day 42. The increase in compressive moduli is attributed to the good compatibility between the alginate matrix and the GO owing to the presence of strong interactions between the alginate macromolecules and the GO nanosheets ^[21]^. After 7 days of cell culture, a slight decrease in the compressive modulus was observed for the 0.5GO, 1GO, and 2GO groups. This result is possibly due to the water uptake ability of GO, decreasing the stiffness of the matrixes. There were no statistically significant differences between the two groups on day 7. Interestingly, a dramatic enhancement of the compressive moduli was observed from day 7 to day 42 of culture in osteogenic media in all groups. There were significant differences within the same group between day 42 and day 1, between day 42 and day 7. The compressive modulus on day 42 was 12 times higher for the 0GO and 0.5GO scaffolds, 7.3 times higher for the 1GO scaffolds and 4.3 times higher for the 2GO scaffolds than on day 7. Marrella et al. ^[17]^ also demonstrated that stiffness of GO/alginate functionalized hydrogels was increased dramatically from one week to four weeks of cell culture. We believe that cell spreading and extracellular matrix mineralization was mostly responsible for the enhanced compressive modulus in later time points. The swelling behavior with excellent water uptake ability of 2GO decreases the compressive moduli of these scaffolds. Compressive stiffness, as well as permeability, is highly correlated with water content: as the water content increases, the material becomes less stiff and more permeable ^[29]^.

### 2.5 Cell viability and spreading

To investigate whether the cell-material interactions are compromised by the presence of GO in the scaffolds, the viability of hMSCs after short- and long-term culture was analyzed. The cytotoxic effects of GO were evaluated by comparing the cell viability results obtained between 0GO and the different GO concentrations. Figure 4A shows representative images depicting cell viability in the 3D bioprinted cell-laden GO scaffolds at day 1, day 7, and day 42 of culture. The predominant green fluorescence evidences the high population of live cells in all groups. Figure 4B shows the quantitative analysis of cell viability in 3D bioprinted cell-laden GO scaffolds on day 1, day 7, and day 42. While all groups were nontoxic (cell viability was above 85%), GO decreased cell death by increasing the concentration of GO, with significant differences detected at 2GO compared with 0GO on day 1 and day 7. Furthermore, all groups supported cell spreading and cell-cell interactions after 42 days of culture. However, GO concentration influenced cell spreading capabilities (Figure 4A and 4C). The cells in the 0GO scaffolds were homogeneously dispersed and showed a spindle-shaped morphology, whereas those in GO scaffolds were larger and more outspread (Figure 4A). Figure 4C shows the influence of GO incorporation on the projected cell area in 3D bioprinted cell-laden scaffolds. The results showed that there were no significant differences within the same group between day 7 and day 1, but significant differences were found within the same group between day 42 and day 1 and between day 42 and day 7. The results indicate that the projected cell area was significantly increased after culture in osteogenic media for 42 days. Meanwhile, the cell area increased with higher GO concentration, and the projected cell area of the GO groups (0.5GO, 1GO and 2GO) was significantly higher than that of the 0GO group on day 7 and day 42. A trend towards a higher projected cell area was observed with increasing GO concentrations; however, no significant difference was found within GO groups. Some researchers have demonstrated that the extraordinary absorption behavior of GO promotes the deposition of nutrition components (such as FBS) on the scaffold surface and is favorable for cell survival and spreading ^[30]^. Our results were similar to the findings of Lee et al. ^[16b]^, who studied the effect of graphene, GO, and polydimethylsiloxane (PDMS) on the 2D differentiation of hMSCs. They found that the cells cultured on graphene and GO exhibited cellular protrusions, which were distinctively different from those cultured on PDMS. The two higher GO concentrations (1GO and 2GO) favored earlier cell spreading at day 7 compared with 0.5GO and 0GO. In addition, higher cell aggregations were visible in the GO groups after 42 days of culture. Su et al. ^[19]^ showed that cells that were encapsulated in GO-incorporated hydrogels (GO concentration of 2 mg/ml) were more metabolically active than those in pure gelatin methacrylate (GelMA) hydrogels.

**Figure 4.**
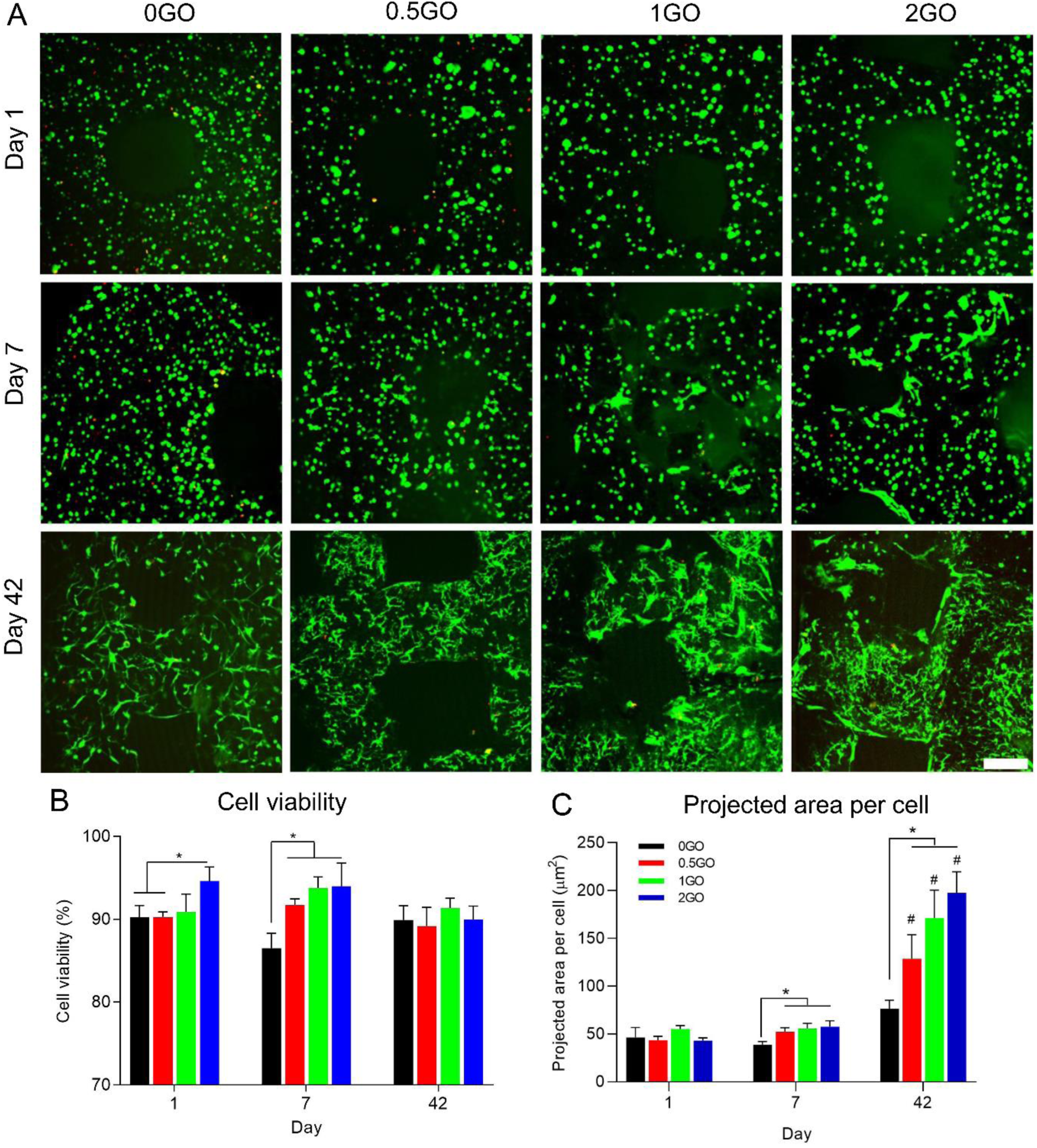
(A) Cell viability in the 3D bioprinted cell-laden GO scaffolds at day 1, day 7 and day 42. Living cells are depicted in green and dead cells in red. The scale bar represents 50 µm. (B) Quantitative analysis of cell viability within 3D bioprinted cell-laden GO scaffolds after 1, 7 and 42 days. (C) Projected cell area in 3D bioprinted cell-laden GO scaffolds cultured for 1, 7 and 42 days in osteogenic media. * P < 0.05, # P < 0.05 within the same group between day 42 and day 1, between day 42 and day 7. Data are shown as mean ± standard deviation (n = 3).

### 2.6 Cell proliferation

Cell proliferation in 3D bioprinted cell-laden GO composite scaffolds was investigated at day 7, 21 and 42 after culture in osteogenic media. GO influenced the DNA content after 7 and 42 days of culture. Figure 5A shows that DNA content was decreased in all groups when cultured in the osteogenic media from day 7 to day 42. Several reasons may contribute to the DNA decrease over time. Some encapsulated cells may have detached from the scaffolds because of the movement of media within the well plate. Previous studies have also reported increased cell death with increased osteogenic differentiation ^[31]^. The extraction of cells may be hampered due to extracellular matrix (ECM) mineralization after 42 days of culture in osteogenic media. However, the DNA content of the 1GO and 2GO groups was significantly higher than that of the 0GO group on day 7 and day 42. No significant difference was detected between the 0GO and GO groups on day 21. This result is consistent with the fact that GO incorporation increases cell proliferation and may be related to the strong adsorption capacity of GO by π-π stacking and electrostatic interactions ^[16b, 16c, 32]^. This adsorptive performance can be considered favorable to FBS adsorption and subsequent cell proliferation. Previous studies ^[33]^ have also reported that the incorporation of carbon-based nanomaterials, such as carbon nanotubes, into ECM-derived substrates support enhanced cellular adhesion and proliferation due to the strong affinity between nanomaterials and ECM proteins.

**Figure 5.**
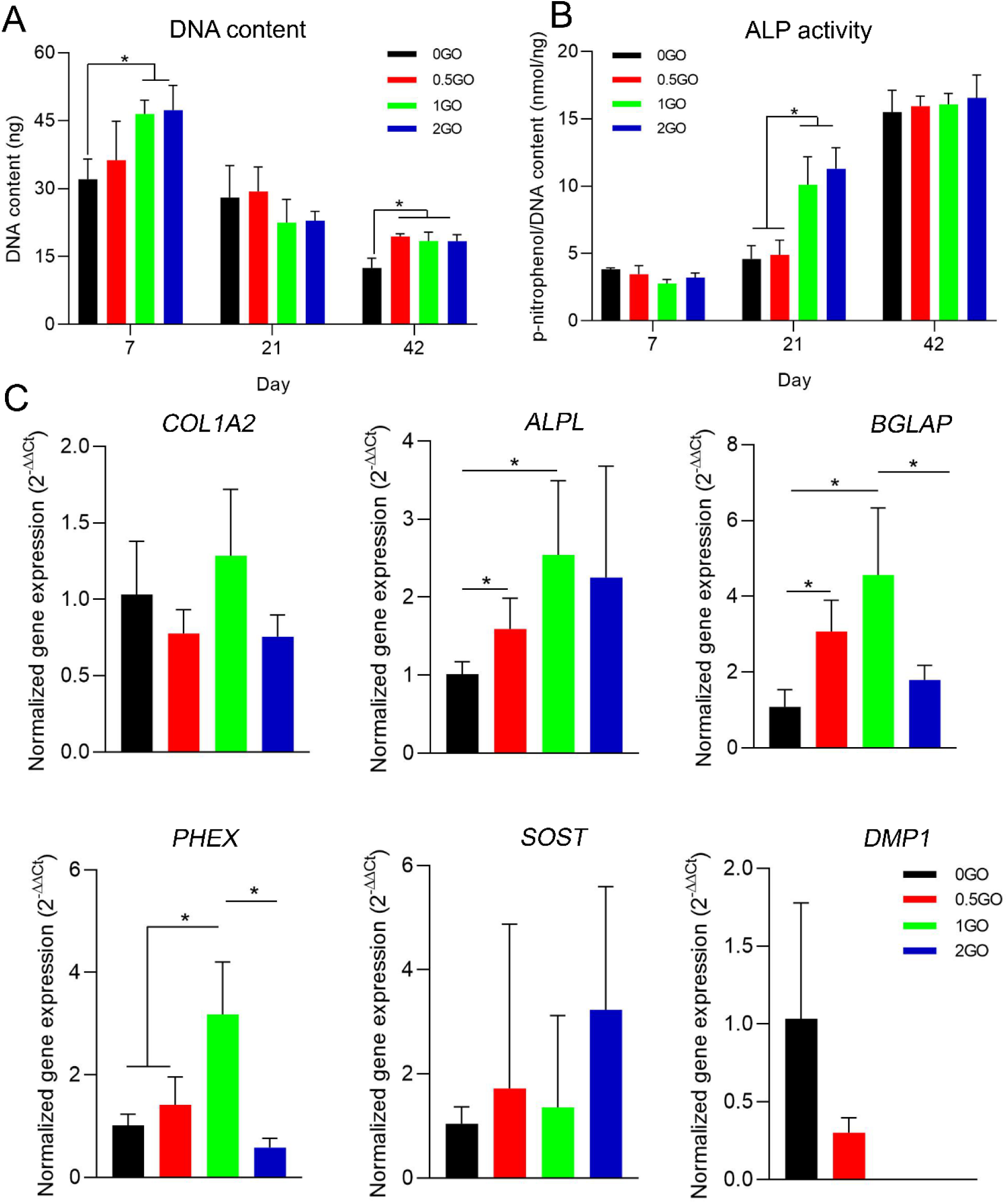
Quantification of the (A) DNA content and (B) ALP activity in the 3D bioprinted cell-laden scaffolds after day 1, 7, 14, 28, and 42 of culture in osteogenic media. (C) Relative gene expression levels of *COL1A2, ALPL, BGLAP, PHEX, SOST* and *DMP1* for the 3D bioprinted cell-laden GO scaffold by RT-qPCR analyses after culturing in osteogenic media for 42 days. * P < 0.05, data are shown as mean ± standard deviation (n = 3).

### 2.7 ALP activity and osteogenic-related gene expression

Alkaline phosphatase (ALP) contributes to the production of ECM for deposition before the initiation of mineralization ^[34]^. ALP activity was measured in the 3D bioprinted cell-laden scaffolds after 7, 14 and 42 days of culture (Figure 5B). In contrast to the results for DNA content, ALP activity increased over time in all groups. There were statistically significant differences between the 0GO and 0.5GO groups and the 1GO and 2GO groups on day 21. No significant difference was detected between the 0GO and GO groups on day 7 or day 42.

The effect of GO on the osteogenic differentiation of hMSCs towards osteoblasts and the osteocyte lineage was further evaluated after culturing in osteogenic media for 42 days (Figure 5 C). Osteoblasts and osteocytes are characterized by a sequence of gene expression. The relative expression of early- (*COL1A2*), middle- (*ALPL*), and late- (*BGLAP*) stage osteoblast-related genes and osteocyte-related genes (*PHEX, DMP1, SOST*) is shown in Figure 5C. *ALPL* and *BGLAP* gene expression was significantly increased in the 0.5GO and 1GO groups compared with the 0GO group. *PHEX* is a gene expressed at the early stage of osteocytes and is involved in regulating skeletal mineralization. In line with the results observed for *ALPL* and *BGLAP, PHEX* gene expression was significantly upregulated in the 0.5GO group compared with the 0GO group and in the 1GO group compared with the 0GO and 0.5GO groups. No differences in *COL1A2* and *SOST* were observed compared with the 0GO group. Although not statistically relevant, the *DMP1* gene was only upregulated in the 0GO and 0.5GO groups. Interestingly, 2GO did not upregulate the expression in any of the genes evaluated compared with 0GO. Our results indicate that appropriate GO incorporation can promote hMSC differentiation into mature osteoblasts and early osteocyte phenotypes, and the optimal group was 1GO. This may be explained by the higher conductivity of GO ^[35]^ and its unique structure with surface oxygenated groups ^[16b, 36]^ compared with alginate or gelatin. It has been shown that the poor conductivity of GO can also have positive effects on cell proliferation and differentiation ^[37]^. Surface oxygenated groups, such as OH^−^ and COO^−^, endow GO with the ability to adsorb specific proteins (present in FBS) ^[30]^ and low-molecular-weight chemicals, such as osteogenic inducers (dexamethasone and β-glycerophosphate) ^[16b]^. Therefore, it is reasonable to speculate that the improved osteogenic differentiation of hMSCs by GO incorporation may be directly related to the specific bioactive groups and extraordinary absorption capacity of GO ^[38]^. However, when the concentration of GO reached 2 mg/ml, it exerted adverse effects on osteogenic cell differentiation. The results are in line with the finding that abundant GO induces oxidative stress in cells and generates reactive oxygen species (ROS) that inhibit cell differentiation ^[39]^.

### 2.8 In situ micro-CT monitoring of mineral formation in 3D bioprinted cell-laden defect scaffolds

We bioprinted 3D cell-laden defect scaffolds with different GO concentrations to mimic *in vivo* calvarial bone defects in mice. 3D cell-laden defect scaffolds were assembled in the bone bioreactors and cultured with osteogenic media for up to 42 days. The influence of GO incorporation on mineral formation in 3D bioprinted cell-laden defect scaffolds was investigated by *in situ* time-lapse micro-CT imaging. Scaffolds were mineralized over time, and 2D mineralized slices could be visualized under micro-CT scanning after culture in osteogenic media for 42 days (Figure 6C). Figure 7A shows the 3D reconstructed time-lapsed micro-CT images in the top view of the 3D bioprinted cell-laden defect scaffolds with different GO concentrations. ECM mineralization increased over time for all groups. On day 7, there was almost no mineral formation in any group. However, after culturing in osteogenic media for 42 days, mineralized 3D defect scaffolds were formed. The defect was kept open, thereby mimicking the *in vivo* critical-size mouse calvarial bone defect model. The defect scaffold shapes were similar to the hydrogel scaffolds after bioprinting, but the overall scaffold area after 42 days of culture was different for the different GO concentrations. The different results of the defect scaffold morphology were in agreement with the scaffold morphology results of the 3D bioprinted cell-laden scaffolds (Figure 3). The 1GO group maintained a better overall area for the defect scaffold morphology than the 0GO group during the 42 days of culture in osteogenic media. Scaffolds in the 0GO group shrank over time, while GO incorporation helped prevent shrinkage and maintain better scaffold fidelity. However, when the GO concentration reached 2 mg/ml, as in the 2GO group, the defect scaffolds swelled and became larger in size than the designed scaffold model after culturing them in osteogenic media for 28 days. GO absorbed the hydrophilic protein and water molecules in a GO concentration-dependent manner, which may have contributed to the swelling behavior of the 2GO defect scaffolds.

**Figure 6.**
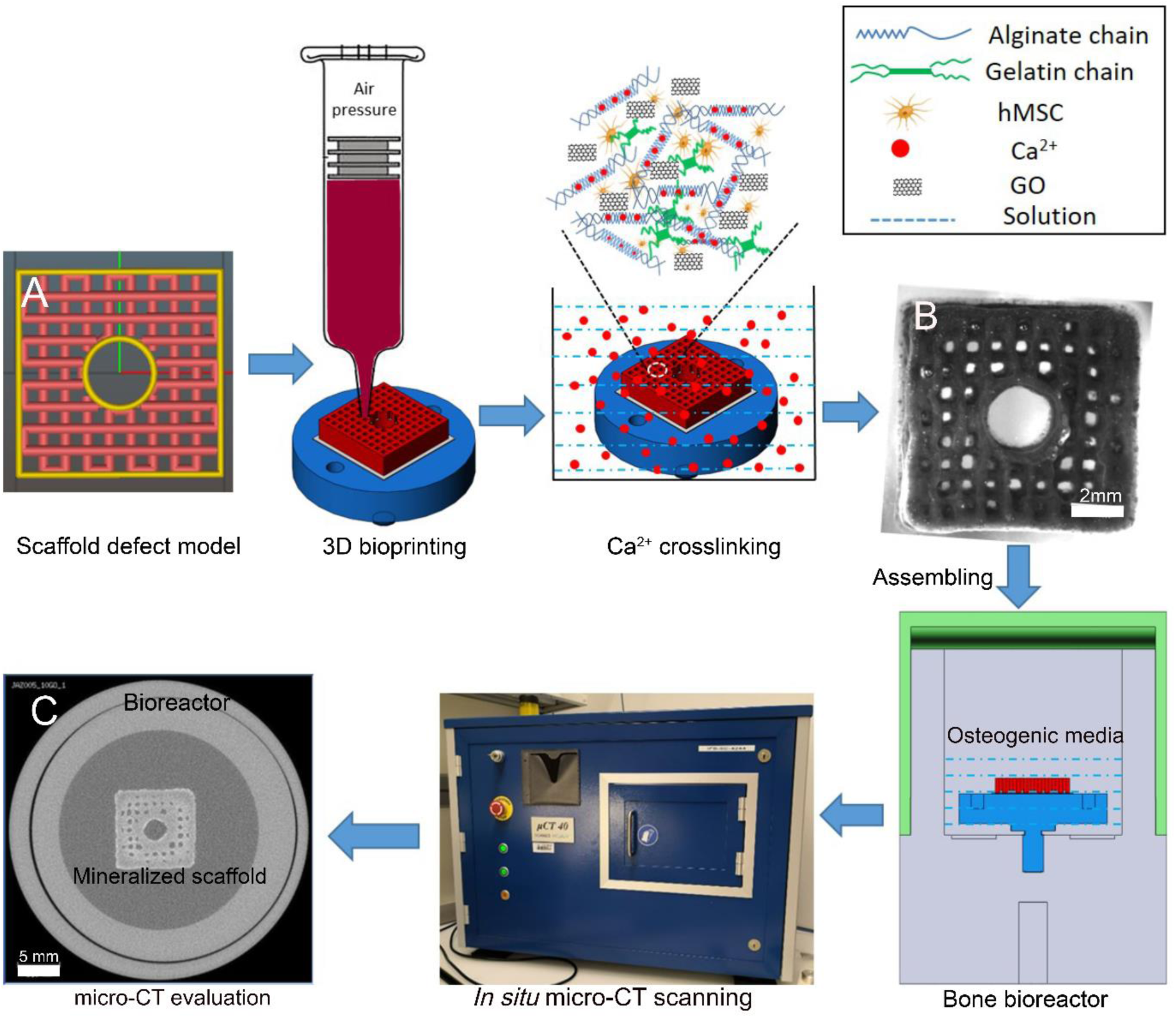
Schematic illustration of 3D bioprinting of the 3D cell-laden defect scaffolds, Ca^2+^ crosslinking, bioreactor assembling, *in situ* micro-CT monitoring and micro-CT evaluation processes. (A) Defect scaffold Stereo Lithography (STL) model, (B) representative images of the 3D cell-laden 2GO defect scaffold at day 1 after printing. (C) 2D CT slide image after culture in osteogenic media for 42 days.

**Figure 7.**
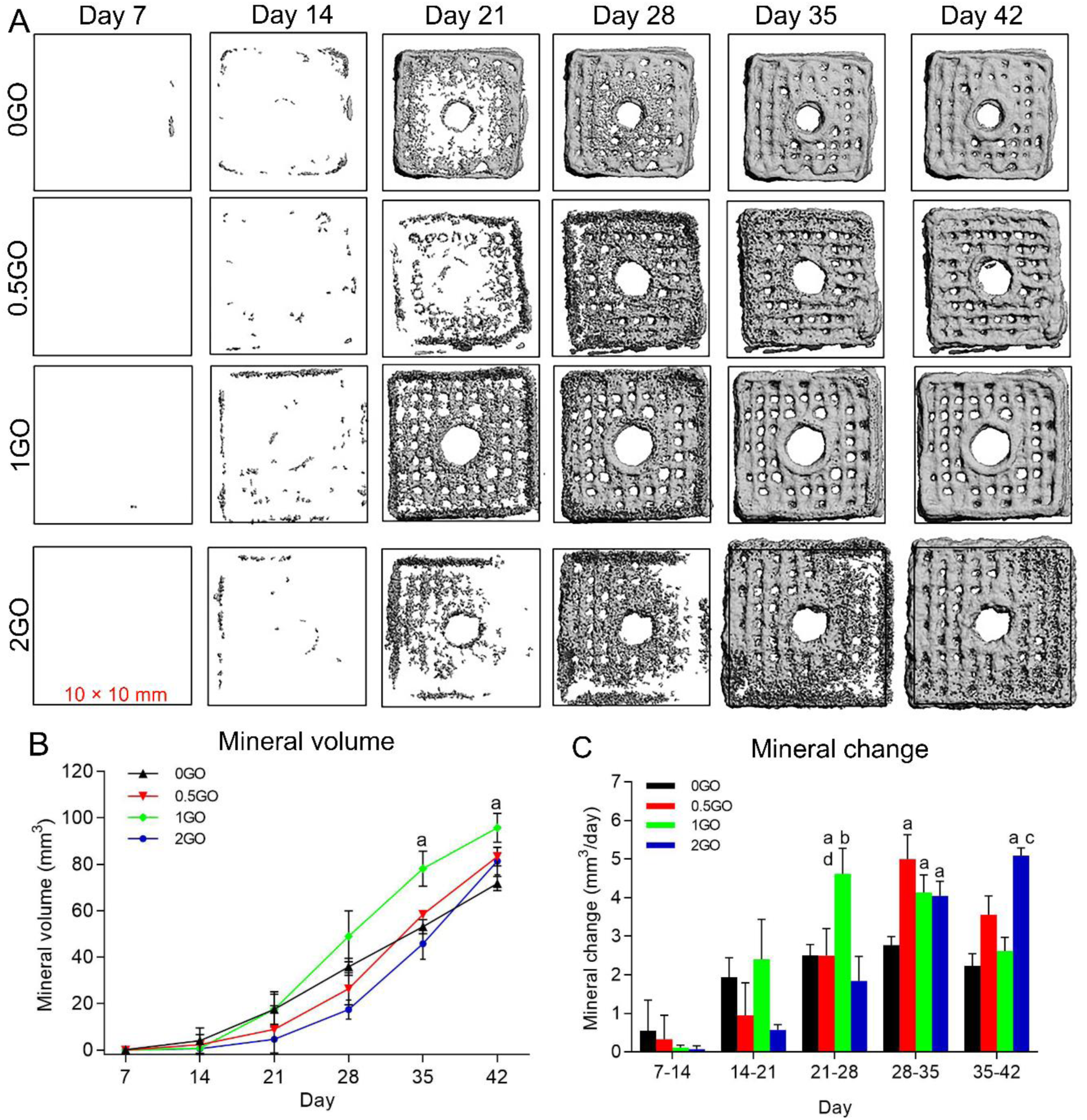
(A) 3D reconstructed time-lapsed micro-CT images at the top view of the 3D bioprinted cell-laden GO scaffolds at each scan timepoint. The size of the square box is 10 × 10 mm, which is the same as the scaffold model. (B) Mineral volume from day 7 to 42. (C) Mineral change between two scans. ^a b c d^ represent significant differences (P < 0.05) compared to 0GO, 0.5GO, 1GO and 2GO group, respectively, within the same time point. Data are shown as mean ± standard deviation (n = 4).

The quantification of mineral volume in 3D bioprinted cell-laden defect scaffolds is shown in Figure 7B. The mineral volume was similar for all groups on day 14 and increased more than ten-fold from day 14 to day 42. The mineral volume for the 1GO scaffolds was 77.4 ± 7.37 mm^3^ on day 35, which was significantly higher than that of the 0GO scaffolds (56.2 ± 1.63 mm^3^). The mineral volumes were 58.5 ± 0.46 mm^3^ and 4.58 ± 5.52 mm^3^ in the 0.5GO and 2GO groups, respectively. On day 42, the mineral volume was 71.8 ± 2.63 mm^3^, 83.3 ± 3.24 mm^3^, 95.7 ± 5.43 mm^3^ and 81.5 ± 4.87 mm^3^ for the 0GO, 0.5GO, 1GO and 2GO scaffolds, respectively. A significant difference in mineral volume was observed between the 1GO and 0GO groups. No significant differences were observed between GO groups on day 42. Furthermore, the mineral changes were calculated every week for all groups (Figure 7C). The period with the highest mineral change varied among groups. The highest mineral change was first achieved on days 21-28 in the 1GO group (4.61 ± 0.57 mm^3^/day), followed by the 0GO group (2.84 ± 0.32 mm^3^/day) and the 0.5GO group (5 ± 0.52 mm^3^/day) on days 28-35. The 2GO group had the highest mineral change of 5.09 ± 0.16 mm^3^/day on days 35-42. The mineral change was significantly higher in the 1GO group than in the other three groups on days 21-28, and the mineral change was significantly higher in the GO groups compared to the 0GO group on days 28-35.

Consistent with the ALP activity and osteogenic-related gene expression results, the GO concentration was chosen for 3D bioprinting of cell-laden alginate/gelatin defect scaffolds, resulting in enhanced osteogenic differentiation and increased ECM mineralization. Lee et al. ^[16b]^ demonstrated that GO can increase the extent of mineralization when hMSCs are cultured on a GO substrate by pre-concentrating osteogenic inducers via π-π stacking. Several studies have shown that GO-modified scaffolds promote new bone formation in bone defects in animal studies ^[38, 40]^. Ma et al. ^[38]^ found that GO-modified β-tricalcium phosphate (GO-TCP) scaffolds significantly promote new bone formation in the bone defects of rabbits compared with pure β-TCP scaffolds. According to these previous studies, it is reasonable to speculate that the improved ECM mineralization of GO groups is closely related to the negative chemical groups, such as COO^−^, for nucleation and crystallization of Ca/P ions ^[41]^. Due to these chemical groups, increasing GO incorporation will change the scaffold surface roughness. Several studies have shown that cells are influenced by surface chemistry and roughness, which may positively affect cell proliferation, differentiation, and mineralization ^[42]^. Mineralized 3D defect scaffolds with embedded human cells have potential applications for use as *ex vivo* bone models that mimic specific human bone tissues. For example, these scaffolds could be used to investigate *ex vivo* the effectiveness of various biomaterials to repair critical-size bone defects, reducing animal usage *in vivo*. They could also be employed to enable more accurate prediction of therapeutic/toxic responses, again reducing animal usage and further decreasing the cost of drug discovery. Furthermore, *in vitro* bone models can be developed by fabricating constructs with diseased or dysfunctional human cells for studying patient-specific tissue pathology or testing new therapeutics in the context of personalized medicine.

### 2.9 Histology staining

ECM mineralization and cell morphology in the 3D bioprinted cell-laden GO defect scaffolds were evaluated by Alizarin red S and H&E staining (Figure 8) after 42 days of culture with osteogenic media in the bone bioreactor. Histological staining was performed from sections at the top region of the 3D cell-laden GO defect scaffolds. Alizarin red S staining was used to qualitatively visualize the highest mineral region (red) and the lower mineral region (yellow). The Alizarin red S staining images showed that sections taken from the 1GO scaffolds exhibited greater red staining intensity, which is an indicator of higher mineral content in the 1GO scaffolds than in the 0GO, 0.5GO and 2GO scaffolds. These results were in line with the mineral volume results of the micro-CT evaluation on day 42. H&E staining showed a uniform cell distribution throughout the whole section for all groups. Cells spread well in all sections. The cells in the 0GO scaffolds were smaller and showed a narrow and elongated morphology, whereas those in the GO scaffolds were larger and displayed polygonal morphology. We also observed, as shown in Figure S1, that the cytoskeleton of the cells was well organized with abundant stress fibers, which are the contractile bundles of actin filaments in the 1GO group, while fewer stress fibers were formed in 0GO scaffolds. Some researchers have shown that cell-matrix junctions formed by integrins when cells were grown on surfaces with higher roughness ^[43]^. The results suggest that GO incorporation into 3D bioprinted cell-laden alginate/gelatin scaffolds changed the scaffold roughness, cell adhesion, and cell morphology with more stress fibers.

**Figure 8.**
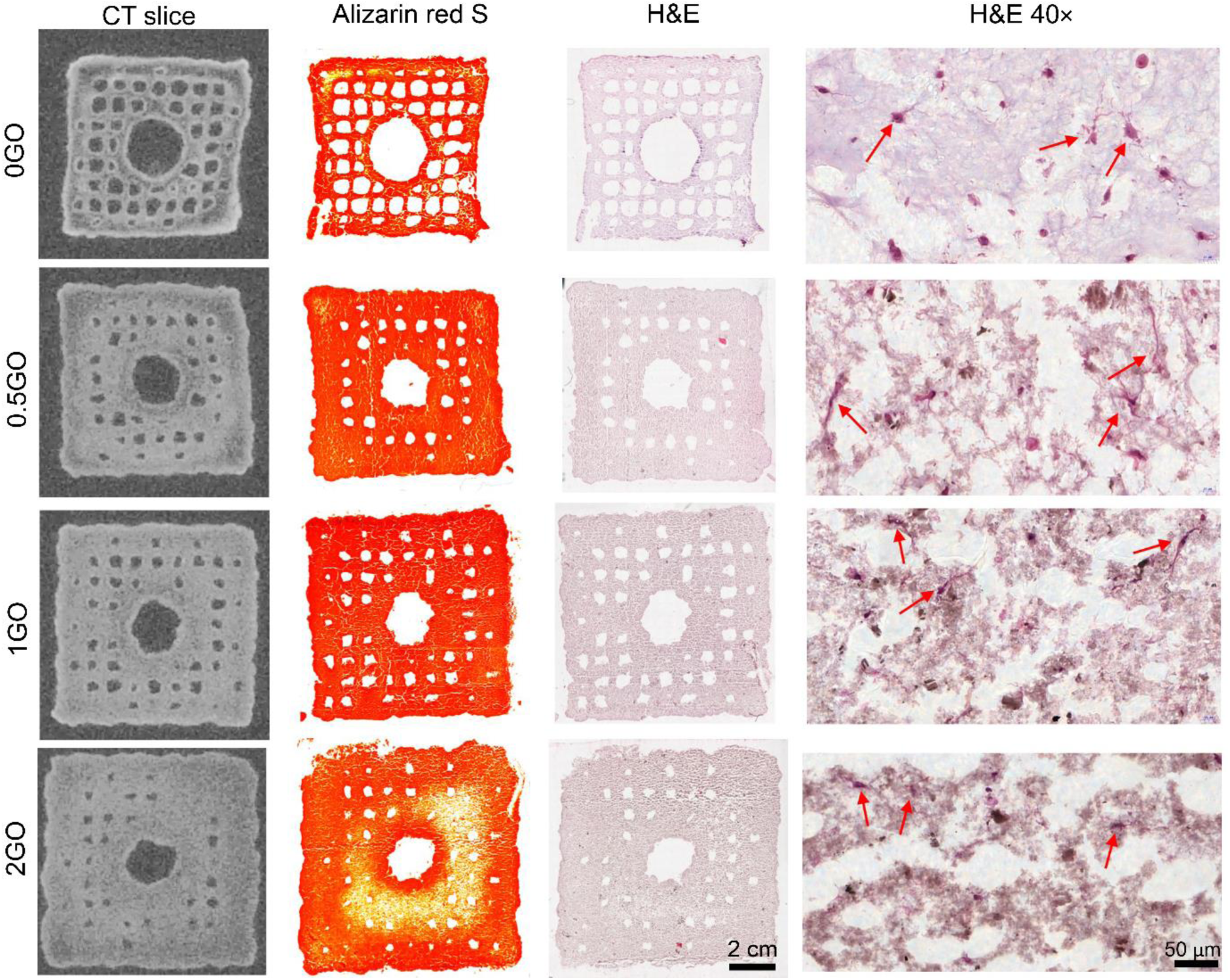
Grayscale CT slice, Alizarin red S staining, H&E staining of the whole section images and 40 × H&E staining images of the 3D bioprinted cell-laden GO scaffolds from the top region after 42 days of culture in the osteogenic media.

## 3. Conclusion

3D cell-laden GO/alginate/gelatin composite bone mimicking scaffolds were successfully fabricated using 3D bioprinting technology. The scaffold fidelity of the GO group was better than that of the 0GO group, and the compressive modulus increased with increasing GO concentration on day 1. After 42 days of culture in the osteogenic media, the 0GO scaffolds shrank, and the 1GO scaffolds maintained the highest-fidelity compared to the scaffolds in the other three groups. The 2GO group scaffolds swelled and had the lowest compressive modulus. Furthermore, GO provided a certain protective effect, improving cell viability on day 1 and day 7. The biocompatibility properties of GO were further manifested by an increase in the overall cell projected area on day 7 and day 42 of scaffolds of all GO concentrations compared with 0GO scaffolds. Nevertheless, the selected GO concentration is highly relevant for promoting osteoblastic/osteocytic cell differentiation and ECM mineralization. The incorporation of GO at 1 mg/ml had the highest osteogenic differentiation by upregulating osteogenic-related gene (*ALPL, BGLAP, PHEX*) expression. Interestingly, GO incorporation increased mineral formation in the 3D bioprinted cell-laden defect scaffolds after 42 days of culture in the bioreactor, which was confirmed by *in situ* micro-CT scanning and histology staining. The 3D bioprinted mineralized defect scaffolds may prove useful as an *ex vivo* bone mimicking model for critical-sized defects, screening of novel compounds or the prediction of toxicity, thereby reducing animal usage. In conclusion, the 3D bioprinted cell-laden scaffolds with 1 mg/ml GO incorporation improved scaffold fidelity and increased osteogenic differentiation and mineralization, and therefore, these scaffolds have extensive applications in bone model fabrication and bone tissue engineering.

## 4. Experimental Section

### Inks preparation and GO characterization

Details of the materials are provided in the supporting information. Four alginate/gelatin/GO composite ink solutions containing different GO concentrations, 0 mg/ml, 0.5 mg/ml, 1 mg/ml and 2 mg/ml GO, were named 0GO, 0.5GO, 1GO and 2GO, respectively. Alginate (70.4 mg, 0.8% w/v) and gelatin (360 mg, 4.1% w/v) were added to a homogeneous glycerol/ phosphate-buffered saline (PBS) (9.1% v/v) solution to prepare the 0GO group as a control, as described previously ^[7]^. The commercial GO product is a 5 mg/ml GO dispersion in water solution. To keep the same alginate and gelation concentrations in the four groups, different PBS concentrations, volumes and GO volumes were used to prepare different GO groups. The amounts of chemicals used for preparing the different alginate/gelatin/GO composite ink solutions are listed in Table 1. As an example, to prepare a 1GO ink solution, 1.6 ml GO solution (5 mg/ml) was mixed with 5.6 ml 1.29× PBS and 0.8 ml glycerol solution to a homogeneous solution. Then, 70.4 mg alginate and 360 mg gelatin were dissolved in the above GO/PBS/glycerol solution at 50 °C in a water bath. The mixture was kept for 12 h to homogenize and fully dissolve the polymer and then pasteurized at 70 °C for 1 h before the next experiments.

**Table 1.** Amounts of chemicals used for GO composite hydrogel preparation.

| Groups | Alginate (mg) | Gelatin (mg) | Glycerol (ml) | PBS (ml) | GO (ml) |
| --- | --- | --- | --- | --- | --- |
| 0GO | 70.4 | 360 | 0.8 | 7.2, 1×PBS | 0 |
| 0.5GO | 70.4 | 360 | 0.8 | 6.4, 1.13×PBS | 0.8 |
| 1GO | 70.4 | 360 | 0.8 | 5.6, 1.29×PBS | 1.6 |
| 2GO | 70.4 | 360 | 0.8 | 4, 1.80×PBS | 3.2 |

The GO morphology was studied by using FEI Talos F200X transmission electron microscopy (TEM) (FEI Company, Hillsboro, USA). The spatial distribution of GO in the GO product and ink solutions was evaluated under a Zoom Stereomicroscope (Leica Microsystems, Heerbrugg, Switzerland).

### Rheology properties

The rheological properties of the bioinks with different graphene oxides (0GO, 0.5GO, 1GO and 2GO) were measured using a HAAKE RheoStress 600 rheometer (Thermo Electron GmbH, Karlsruhe, Germany). All measurements were performed in rotation with a cone-plate geometry with a diameter of 35 mm, a cone-to-plate distance of 105 µm and a cone angle of 2°. A liquid sample of 400 µl bioink was placed on the plate at 37 °C. Viscosity and shear stress were measured at 37 °C with a rotatory test setup by varying the shear stress from 0.01 s^−1^ to 100 s^−1^. Shear recovery was performed under oscillation by measuring the storage modulus (G’) and the loss modulus (G’’) at a frequency of 5 rad s^−1^ and a low strain of 0.5% for 60 s. The strain was then increased to 500% at the same frequency for 60 s before returning to 0.5% strain for 240 s. The thermoresponsive properties were analyzed by G’ and G’’, which were assessed by oscillatory measurements at a frequency of 1 rad/s and 1% strain. Before rheology measurements, the hydrogels were warmed up to 37 °C. For the measurements, the hydrogels were cooled from 30 °C to 10 °C at a rate of 0.5 °C min^−1^. The gelation temperature was taken as the temperature when G’ and G’’ were equal. All measurements were repeated three times to ensure consistency of the obtained results.

### 3D bioprinting of acellular GO scaffolds

In this study, a microextrusion-based two-syringe cell bioprinter (INKREDIBLE^+^, CELLINK, USA) was used to print the different alginate composite scaffolds. The scaffold model used was a lattice-rod structure (10 mm × 10 mm × 2.5 mm), which was converted into a G-Code file using Slic3r with 0.17 mm layer height (15 layers), 0.76 mm pore size and 0.32 mm nozzle diameter. The ink solution was prepared by thoroughly mixing 1 ml of hydrogel solution with 100 µl DMEM with magnetic fish at 200 rpm/min for 1 minute at room temperature (20 °C). At room temperature, the ink solution is in a liquid state and cannot be extruded as a stable filament. To allow printability, the ink was introduced into a polyethylene injection cartridge (5 cc) and cooled in a Styrofoam box containing ice for 8 min. Reducing the temperature of the ink quickly and gaining fast gelation conferred the thermosensitivity property of the gelatin. The cartridge was fixed onto the printing device, and the G-Code model scaffold design was loaded into the printer’s disk. Scaffolds were printed layer-by-layer via ink extrusion as fibers to create 3D scaffolds on the bioprinter platform surrounded with dry ice. Typically, air pressure was set to 50 kPa, speed was 2 mm/s, and nozzle size was 27 G. Immediately after printing each scaffold, the scaffolds were crosslinked with 2% (w/v) CaCl2 solution for 10 min to maintain structural integrity for long-term stability. To remove the excess CaCl2, the scaffold was washed 3 times in cell culture media, and each wash was 10 seconds. Scaffolds were cultured in osteogenic media (DMEM, 10% FBS, 1% P/S/F, 50 µg/ml AA, 100 nM Dex, 10 mM β-GP) in an incubator (37 °C, 5% CO2). The osteogenic media was changed every two days.

### 3D bioprinting of cell-laden GO scaffolds

hMSC (Lonza, Walkersville, MD, USA) were isolated from bone marrow aspirates and characterized as described previously [44]. P3 hMSC were cultured in expansion media (DMEM, 10% FBS, 1% P/S/F, 1% NEAA and 1 ng/ml bFGF) in an incubator (37 °C, 5% CO2) for 7 days. All cells were cultured in tissue culture polystyrene triple flasks, and cells at passage 4 were used for the subsequent experiments. After harvesting, the cells were resuspended at a density of 5 million cells in 100 µl cell culture media (DMEM, 10% FBS, 1% P/S/F). The bioink was prepared by mixing the 100 µl cell suspension with a 1 ml ink solution. The scaffold model, 3D bioprinting, crosslinking and scaffold culturing processes were the same as described above for 3D printing of acellular scaffolds.

### Scaffold structure

The influence of GO concentration on the structure of the 3D bioprinted scaffolds over time was observed using a Zoom stereomicroscope (Leica Microsystems, Heerbrugg, Switzerland). Two-dimensional top view images of the 3D bioprinted acellular scaffolds (n = 3) were captured after incubation of the scaffold for 7 days in osteogenic media and under standard cell culture conditions (37 °C and 5% CO_2_). The morphological changes of the 3D bioprinted cell-laden scaffolds (n = 3) were observed on day 1, 7 and 42 of cell culture in osteogenic media. All images were taken at 1x magnification. Scaffold structure parameters, including pore size, filament diameter, pore area, and scaffold overall area, were manually measured and analyzed using ImageJ (National Institutes of Health, USA). The scaffold morphology parameters of the 3D bioprinted scaffolds on day 0 used the same parameters as the scaffold model. Pore size, filament diameter, and pore area were measured in at least ten different positions for each scaffold and averaged for a final value. The scaffold overall area was measured in three scaffolds per group and then averaged.

### Cell viability

Cell viability in the 3D bioprinted cell-laden GO scaffolds was assessed using a LIVE/DEAD® Viability/Cytotoxicity assay after 1, 7 and 42 days of culture in osteogenic media. After washing twice in PBS, samples were transferred to a 2 μM Calcein-AM and 4 μM ethidium homodimer (EthD-1) solution and cultured in a CO_2_ cell incubator for 40 min. Then, the cell-laden scaffolds were thoroughly rinsed with PBS twice and kept on sterile chambered cover glasses (Lab-Tek™, Thermo Scientific) with 1.5 ml cell culture media. Live cells emitted green fluorescent light and were visualized under the green fluorescent protein (GFP) channel (emission: 525 nm). Dead cells emitted red fluorescent light and were visualized under the mCherry fluorescent protein channel (emission: 630 nm). Confocal microscopy (Visitron Spinning Disk, Nikon Eclipse T1) was used to visualize live and dead cells, and six representative images of each scaffold were captured (magnification of 10x) for this purpose. Both live and dead cell numbers were manually counted and analyzed using ImageJ (National Institutes of Health, USA). Cell viability was calculated as the percentage of live cells as a fraction of the total cell count.

### Scaffold mechanics

The mechanical properties were characterized by measuring the compressive modulus, which was assessed on a Zwick material testing machine (Zwick 1456, Ulm, Germany) with a 10 N load cell at room temperature. An unconfined uniaxial compression test was performed under displacement control, with a preload of 5 mN and a strain rate of 1 min^−1^ until 60% maximal deformation of the construct was reached. Acellular scaffolds were tested after culture in osteogenic media for 1, 7 and 42 days. The compressive modulus was calculated from the linear region of the stress-strain curve. Three scaffolds (n = 3) were measured per group, and the average value and standard deviation were obtained.

### Cell proliferation and ALP activity

After culturing in osteogenic media for 7, 21 and 42 days, scaffolds were washed twice in PBS. Mini Beadbeater TM (Biospec, USA) was used for scaffold disintegration and cell lysate. The scaffolds were disintegrated in 0.45 ml of 0.2% (v/v) Triton X-100 and 5 mM MgCl2 solution using two steel beads and the minibead beater. Each sample was processed three at 25 000 RPM for 10 seconds each time, and the sample was placed on ice between cycles for cooling. Cell proliferation was determined by measuring the DNA content obtained from the cell lysates. After 48 h incubation at room temperature at which time scaffolds had been disintegrated, the precipitate was separated by centrifugation at 3000 g for 10 min at room temperature. A Quant-iT ™ PicoGreen assay (Life Technologies, Switzerland) of the supernatant fraction was performed according to the manufacturer’s instructions. The DNA content was calculated with three samples in every group and averaged. ALP activity measurements were carried out directly after scaffolds had been disintegrated. Eighty microliters of the supernatant were mixed with 20 µL of 0.75 M 2-amino-2-methyl-1-propanol (abcr GmbH, Germany) buffer and 100 µL 100 mM p-nitrophenyl phosphate solution (Honeywell Fluka(tm), Hungary) and incubated for 2 min in a 96-well plate. Then, 0.2 M NaOH was added to stop the conversion of p-nitrophenyl phosphate to p-nitrophenol. The absorbance values were measured at 405 nm. ALP activity was calculated using a p-nitrophenol standard curve included in the assay. The calculated ALP activity was then normalized by the DNA content measured for each scaffold.

### Osteogenic-related gene expression

Total RNA of the 3D cell-laden scaffolds was extracted using TRIzol Reagent (Invitrogen(tm), Thomas Fisher Scientific, USA), pellet pestles (Sigma-Aldrich, USA) and an RNeasy Micro Kit (Qiagen, Switzerland) according to the manufacturer’s instructions. The RNA isolation method was performed as previously described ^[45]^. RNA purity and concentration were quantified at 260 nm using a NanoDrop 2000 (Thermo Fisher Scientific, USA). Total RNA (410 ng) was reverse transcribed into cDNA using an iScript™ Synthesis Kit on a T100™ thermal cycler (BIO-RAD, USA) according to the manufacturer’s protocol. At the end of the procedure, cDNA samples were stored at −20 °C until further analysis. Quantitative real-time PCR (RT-qPCR) was performed in a CFX96TM real-time PCR system (BIO-RAD, USA) using a probe detection method. TaqMan Fast Universal PCR Master Mix (Thermo Fisher Scientific, USA) was used. Each gene was run in duplicate in a single plate, which included all the samples as well as controls. PCR conditions were as follows: denaturation (95 °C for 20 s) followed by 43 amplification cycles (95 °C for 1 s, 60 °C for 20 s). RT-qPCR was performed for six of the target genes (Alkaline phosphatase (*ALPL*), Collagen type I alpha 2 chain (*COL1A2*), Bone gamma-carboxyglutamate protein (*BGLAP*), Phosphate regulating endopeptidase homolog, X-linked (*PHEX*), Sclerostin (*SOST*), Dentin matrix acidic phosphoprotein 1 (*DMP1*)) and one housekeeping gene (Glyceraldehyde-3-phosphate dehydrogenase (*GAPDH*)). Details of the primers were provided in Table S1. Ct values obtained for each sample were normalized to the housekeeping gene. Data were analyzed using the comparative Ct method (2^−ΔΔCt^) and presented as a fold-change in gene expression of the GO groups versus the 0GO group.

### In situ micro-CT monitoring of mineral formation in 3D bioprinted cell-laden defect scaffolds

SolidWorks software (Dassault Systemes S.A, Paris, France) was used to design a square model (10 mm × 10 mm × 2.5 mm) containing a single circle defect of 3 mm in diameter in the center and saved as a stereolithography file (STL). The scaffold defect model was converted into a G-Code file using Slic3r with 0.17 mm layer height, 0.76 mm pore size and 0.32 mm nozzle diameter. The bioprinting processes were the same as described above for printing the lattice-rod scaffolds. To keep the scaffolds stable such that they could not move during micro-CT scanning, scaffolds were directly bioprinted on a sterile 3M two-sided tape (Scotch, 3M, USA) that was adhered to the bioreactor platform, as previously described [45]. The scaffold defect model, 3D bioprinting method, Ca^2+^ crosslinking, bioreactor assembly and *in situ* micro-CT scanning and micro-CT evaluation processes are shown in Figure 6.

*In situ* micro-CT images of all defect scaffolds cultured in bone bioreactors with 5 ml osteogenic media were captured at 7, 14, 21, 28, 35 and 42 days to monitor mineral formation, as described previously ^[46]^. All bioreactors were scanned using a micro-CT 40 (SCANCO Medical AG, Brüttisellen, Switzerland) at a voxel resolution of 36 µm. The energy was set to 45 kVp, the intensity was 177 µA, the integration time was 200 ms, and two-fold frame averaging was chosen. The reconstructed images were Gaussian filtered with a filter width of 1.2 and support of 1 to suppress noise. Mineralized tissue was segmented by thresholding at a grayscale value of 145 (corresponding to a mineral density of 83.44 mg HA cm^−3^) ^[45]^. Unconnected particles smaller than 50 voxels were removed using component labeling. The resulting 3D volume was evaluated morphometrically for mineral volume (mm^3^) and mineral change (mm^3^/day) ((mineral change = mineral volume day m – mineral volume day n)/(m-n)), as described previously ^[46-47]^.

### Histology staining

After 42 days of culture in osteogenic media, the scaffolds were washed twice with PBS and fixed in 4% formaldehyde in 10 mM CaCl2 and 0.15 M NaCl solution for 2 hours. Then, the samples were placed in a 30% sucrose solution with 10 mM CaCl2 solution for 16 hours at room temperature. After that, the samples were cryoembedded using optimum cutting temperature compound (OCT, VWR) and snap-frozen on dry ice. The samples were sectioned using Kawamoto’s cryofilm type 2C (SECTION-LAB Co. Ltd., Japan) to 7 µm thickness using a cryotome (CryoStar NX70, Thermo Scientific) according to the Kawamoto protocol ^[48]^. Before staining, the sections were fixed on microscope slides using 1% chitosan adhesive. For this purpose, two drops of chitosan solution were deposited on the slide, and the sections were placed on the slide and kept in a running fume hood to allow the chitosan to dry ^[48]^. Alizarin red S staining was performed to visualize mineralized ECM, and hematoxylin and eosin (H&E) staining was performed to visualize cell nuclei and dendrites. The slides were imaged on an automated slide scanner (Pannoramic 250 Flash II, 3Dhistech, Hungary) using the brightfield channel with a magnification of 20x. Alizarin red S staining to identify the mineral location and was compared with the micro-CT slice, as shown in our previous work [45].

### Statistical analysis

GraphPad Prism 8 (GraphPad, California, USA) was used to perform the statistical analysis of the obtained data. Groups were compared at one timepoint using one-way ANOVA together with pairwise comparisons, followed by Tukey corrections. The 0GO, 0.5GO, 1GO and 2GO groups were compared at different timepoints using two-way ANOVA together with Tukey’s multiple comparisons test. * P < 0.05 was considered statistically significant, and ^a b c d^ represent significant differences (P < 0.05) compared with the 0GO, 0.5GO, 1GO and 2GO groups, respectively, at the same time point. Data are shown as mean ± standard deviation (n = 4).

## Acknowledgments

J. Zhang gratefully acknowledges the financial support from the Chinese Scholarship Council (CSC, No.201508310116). The authors thank Dr. H. Huang from the Laboratory for Multifunctional Materials in ETH Zurich for kindly providing the GO product. The authors also thank the Scientific Center for Optical and Electron Microscopy (ScopeM) of ETH Zurich for providing the microscopy facilities and the Tissue Engineering and Biofabrication group in ETH Zurich for providing the INKREDIBLE^+^ bioprinter.

