## supplemental for "3D Bioprinting of Graphene Oxide-Incorporated Cell-laden Bone Mimicking Scaffolds for Promoting Scaffold Fidelity, Osteogenic Differentiation and Mineralization": Supporting information.pdf

*Jianhua Zhang, Hande Eyisoylu, Xiao-Hua Qin, Marina Rubert and Ralph Müller**

Institute for Biomechanics, ETH Zurich, Leopold-Ruzicka-Weg 4, 8093 Zurich, Switzerland

**Experimental Section**

*Materials*: Alginic acid sodium salt (Mw = 10,000-600,000) was purchased from AppliChem GmbH (Darmstadt, Germany). Type A gelatin from porcine skin (Mw: 50,000-100,000), dexamethasone (Dex), ascorbic-acid (AA) and beta-glycerophosphase (β-GP) were purchased from Sigma-Aldrich (Buchs, Switzerland). Glycerol (99.5%) was purchased from Hänseler AG (Herisau, Switzerland). Phosphate buffered saline (PBS) tablets were purchased from Medicago (Uppsala, Sweden). Calcium chloride ($CaCl_2$) was purchased from VWR International (Dietikon, Switzerland). Dulbecco's modified Eagle's media (DMEM), fetal bovine serum (FBS), penicillin-streptomycin-fungizone (P/S/F), non-essential amino acids (NEAA) and fibroblast growth factor-basic (bFGF) were purchased from Life Technologies (Zug, Switzerland). The target osteogenic related genes ((Alkaline phosphatase (*ALPL*, Hs01029144_m1), Collagen type I alpha 2 chain (*COL1A2*, Hs01028956_m1), Bone gamma-carboxyglutamate protein (*BGLAP*, Hs01587814_g1), Phosphate regulating endopeptidase homolog, X-linked (*PHEX*, Hs01011689_m1), Dentin matrix acidic phosphoprotein 1 (*DMP1*, Hs01009391_g1) and Sclerostin (*SOST*, Hs00228830_m1)) and housekeeping gene (Glyceraldehyde-3-phosphate dehydrogenase (*GAPDH*, Hs02758991_g1) was purchased from TaqMan Assays (Thermo Fisher Scientific, USA). Phalloidin (Alexa-Fluor 555, Invitrogen) was purchased from Invitrogen and Hoechst 33342 was purchased from Thermo Fisher Scientific. 5 mg/ml graphene oxide (GO) dispersion in water solution was purchased from RoyalElite RoyCarbon (Shanghai, China)

*Actin staining*: After 42 days of culture in osteogenic media, the 3D bioprinted cell-laden scaffolds (0GO and 1GO) were fixed in 4% formaldehyde in 10 mM CaCl2 and 0.15 M NaCl solution for 2 h. The scaffolds were blocked in 0.1% bovine serum albumin (BSA) and 0.3% Triton/PBS solution for 30 min. The whole constructs were stained with phalloidin (1:100) and Hoechst (1:200) for 40 min to visualize filamentous F-actin and cell nuclei, respectively. The scaffolds were visualized with a confocal microscope (Confocal Zeiss LSM 880 airyscan, Jena, Germany).

Table S1. Primers sequences used for the analysis of relative gene expression by RT-qPCR.

| Gene symbol | Gene name | Assay ID | Amplicon Length |
|---|---|---|---|
| *COL1A2* | Collagen type I alpha 2 chain | Hs01028956_m1 | 71 |
| *ALPL* | Alkaline phosphatase, liver/bone /kidney | Hs01029144_m1 | 79 |
| *BGLAP* | Bone gamma-carboxyglutamate protein | Hs01587814_g1 | 138 |
| *PHEX* | Phosphate regulating endopeptidase homolog, X-linked | Hs01011689_m1 | 91 |
| *SOST* | Sclerostin | Hs00228830_m1 | 81 |
| *DMP1* | Dentin matrix acidic phosphoprotein 1 | Hs01009391_g1 | 106 |
| *GAPDH* | Glyceraldehyde-3-phosphate dehydrogenase | Hs02758991_g1 | 93 |

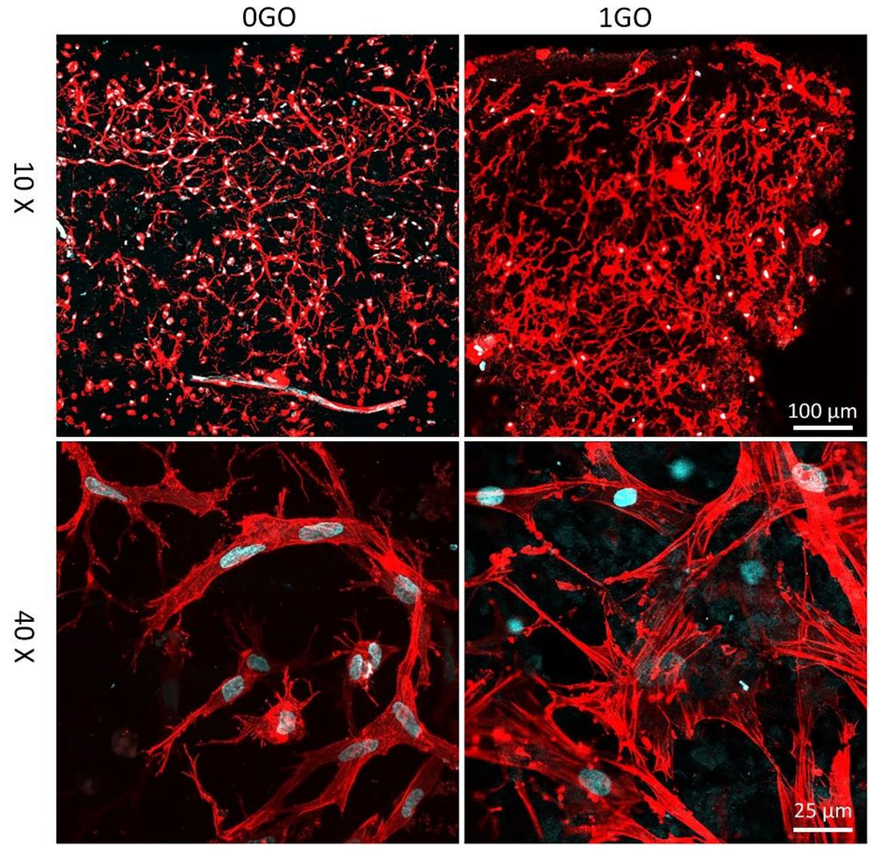


Figure S1. Actin staining images of 3D bioprinted cell-laden 0GO and 1GO scaffolds on day 42 after culture in osteogenic media under 10× and 40× magnification.
